# Group-level patterns emerge from individual speed as revealed by an extremely social robotic fish

**DOI:** 10.1101/2020.06.10.143883

**Authors:** Jolle W. Jolles, Nils Weimar, Tim Landgraf, Pawel Romanczuk, Jens Krause, David Bierbach

**Author notes:** Correspondence should be addressed to: Jolle W. Jolles and David Bierbach.

## Abstract

Understanding the emergence of collective behaviour has long been a key research focus in the natural sciences. Besides the fundamental role of social interaction rules, a combination of theoretical and empirical work indicates individual speed may be a key process that drives the collective behaviour of animal groups. Socially-induced changes in speed by interacting animals make it difficult to isolate the effects of individual speed on group-level behaviours. Here we tackled this issue by pairing guppies with a biomimetic robot. We used a closed-loop tracking and feedback system to let a robotic fish naturally interact with a live partner in real time, and programmed it to strongly copy and follow its partner’s movements while lacking any preferred movement speed or directionality of its own. We show that individual differences in guppies’ movement speed were highly repeatable and shaped key collective patterns: higher individual speeds resulted in stronger leadership, lower cohesion, higher alignment, and better temporal coordination in the pairs. By combining the strengths of individual-based models and observational work with state-of-the-art robotics, we provide novel evidence that individual speed is a key, fundamental process in the emergence of collective behaviour.

## Introduction

Understanding the emergence of collective behavioural patterns has long been a key research focus in the natural sciences. Considerable theoretical and experimental work has accumulated that describes how complex collective patterns may arise via relatively simple mechanisms [1,2], including the role of phenotypic heterogeneity within and among groups [3]. A fundamental insight is that social interaction rules at the individual level – such as avoiding others that are too near and approaching those far away – can explain the large-scale cohesion, coordination, and decision-making of animal groups [2,4].

Most animals control their motion by modulating their speed and turning, and this speed regulation has been shown to be crucial for the attraction and avoidance behaviour when animals group and interact [5–7]. Hence, individual speed may be an additional fundamental factor that underlies the emergence of the global properties of groups. Indeed, both short-term changes and heterogeneity in speed have been linked to a range of group-level properties, such as group cohesion, structure, shape, coordination, and leadership by both theoretical analyses [8], simulations [1,4,9], and empirical work [6,10–14]. Importantly, grouping individuals may differ in their preferred and optimal movement speeds yet must also coordinate and adjust their behaviour to successfully group together [3,15]. Such socially-induced changes in speed by individuals interacting with one another make it difficult to isolate the effects of individual speed for group-level properties. While with agent-based simulations one can separately model such effects and thereby make important predictions for collective behaviour, they are no substitute for empirical data of real animal groups [2].

Recent advances in the field of robotics now make it possible to combine the strengths of agent-based models and behavioural experiments, with robotic individuals behaving like realistic-looking conspecifics and interacting naturally with live animals [16–18]. Here we present results from experiments using live guppies (*Poecilia reticulata*) that were swimming with an interactive biomimetic fish-like robot (‘robofish’) to examine the role of animal’s individual movement speed on collective behavioural patterns. We combined high-definition video tracking and closed-loop feedback system that used interaction rules from well-known agent-based models [4] to steer the robot interactively in real time [19,20]. By programming the robot to always follow its partner and copy its behaviour, while excluding any preferred swimming speed or directionality, we were able to determine how group-level properties in terms of leadership, cohesion, alignment, and temporal coordination emerged from individual differences in guppies’ movement speed.

## Methods

We used lab-reared descendants of wild-caught Trinidadian guppies that were housed in large, randomly out-bred mixed-sex stock tanks under controlled laboratory conditions (12h:12h light:dark; 26°C). We randomly selected 20 naïve adult females (standard length ‘BL’: 31.7 ± 0.8 mm) and moved them to individual holding tanks (40 ⅹ 20 × 25 cm). The following week, we tested fish first without robofish to assess their preferred movement speed (wk 2) and then twice with the robofish (wk 3; trial 2 five days later). Throughout, fish were fed twice daily *ad libitum* with TetraMin flake food.

The test arena consisted of a large white glass tank (88 cm × 88 cm, water height 7.5 cm) that was illuminated from above and enclosed to minimize potential external disturbances. Fish were moved from their individual holding compartment to the experimental tank where they were allowed to acclimatize in an opaque PVC cylinder in the corner of the tank. After one minute the cylinder was raised, the fish filmed from above for 10min, and its movements automatically tracked at 30fps using BioTracker [21]. For the trials with the robotic fish we used a 3D-printed fish replica resembling a female guppy that was connected via magnets to a two-wheeled robot below the tank (see Figure S1 and for details [20]). The robot was controlled via a closed-loop system whereby the movements of the fish were tracked and fed-back to the robot control. The robot unit then adjusted its position and orientation in real-time (i.e. with 30 herz) to result in natural response times. Robofish was circling in front of the acclimatisation cylinder and as soon as the guppy was released from the cylinder started its interactive behaviour.

Robofish’s interactive behaviour was based on the zonal model [4] and allowed the robot to copy the live fish’s motions and follow at a similar speed without any own speed or directional preference (see Figures 1 and S2). We programmed robofish to orientate towards the live fish’s position and maintain a distance of 10 - 15 cm (∼ 4 BL, ‘optimal distance zone’), reflecting observations of wild guppies [22]. This resulted in robofish following at the instantaneous speed exhibited by the live fish while it was in this optimal distance zone. The robofish gradually decreased or increased its speed when the focal fish got into the graduation zone (3 - 10 cm) or beyond the optimal distance zone respectively. If the focal fish was at a less than 3 cm away, robofish stopped moving forward but kept turning at its location to focus on the live fish’s position. The maximum speed and acceleration of robofish were set to reflect that observed for the guppies when alone (25 cm/s and 2.5cm/s^2^ respectively, see Figure S3), with its max. turning rate >360°/s.

**Figure 1.**
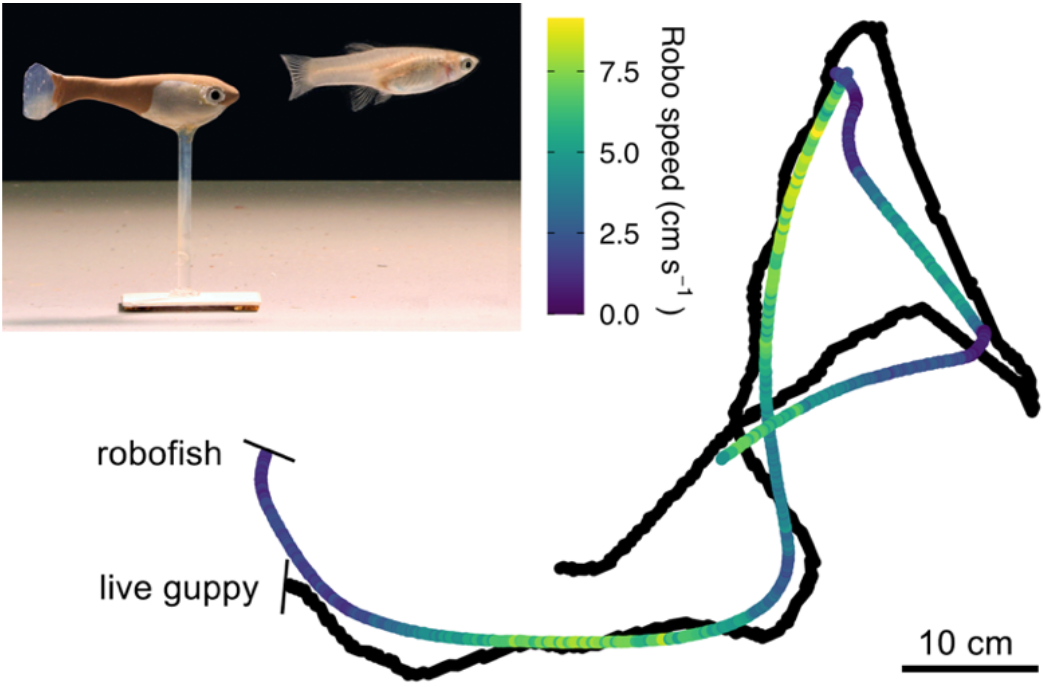
Tracking data (∼ 1min) of a randomly selected pair with the speed of the robofish coloured blue (low) to yellow (high), showing how it followed the position and movements of its partner by natural changes in speed (see further Figure S2). Inset shows a photo of robofish following a guppy.

Tracking data was checked for errors, processed to correct for missing frames, and converted to mm. Subsequently, based on the centroid of each individual (focal and robotic) we calculated speed and heading as well as inter-individual distance. For each trial we computed fish’s median speed, the median inter-individual distance, median difference in heading angle, and proportion of time the focal fish was in front (when moving >0.5cm/sec). In addition, we computed their coordination by running temporal correlations of both individuals’ change in speed and heading, with a higher correlation indicating movement changes were better copied between them. We used a linear-mixed modelling approach to investigate relationships in the individual- and group-level metrics as well as repeatability in behaviour. Further details of our methods and statistics can be found in the Electronic Supplementary Material. All reported experiments comply with the current German law approved by LaGeSo (G0117/16 to Dr. D. Bierbach).

## 3. Results

There were large and significant among-individual differences in guppies’ movement speed across the two robofish trials (*R* = 0.70, 95% confidence interval (CI) = 0.37 - 0.88), which correlated well with their movement speed when tested alone (*χ*^2^ = 9.70, *p* = 0.002, R^2^_mar_ = 0.32, Figure S4). Although the majority of fish slowed down when tested with robofish (30/38 trials), there was large among-individual variation in fish’s speed relative to that expressed in the solo assay (0.62 ± 0.06, 95% CI = 0.15 - 1.29; 1 would indicate no change in speed), which was not linked to their solo speed (*χ*^2^ = 0.155, *p* = 0.212, R^2^_mar_ = 0.06).

Robofish conformed extremely well with the speed of their guppy partner (*χ*^2^ = 87.97, *p* < 0.001, R^2^_mar_ = 0.92, Figure S5) and consistently exhibited a slightly slower speed (speed difference: −0.24 ± 0.03 cm/s). As a consequence, robofish primarily occupied the following position (38/38 trials, Figure S6), which strongly increased with the speed of the focal fish (*χ*^2^ = 40.77, *p* < 0.001, R^2^_mar_ = 0.68), with the fastest guppies leading more than 90% of the time (Figure 2a,b).

**Figure 2.**
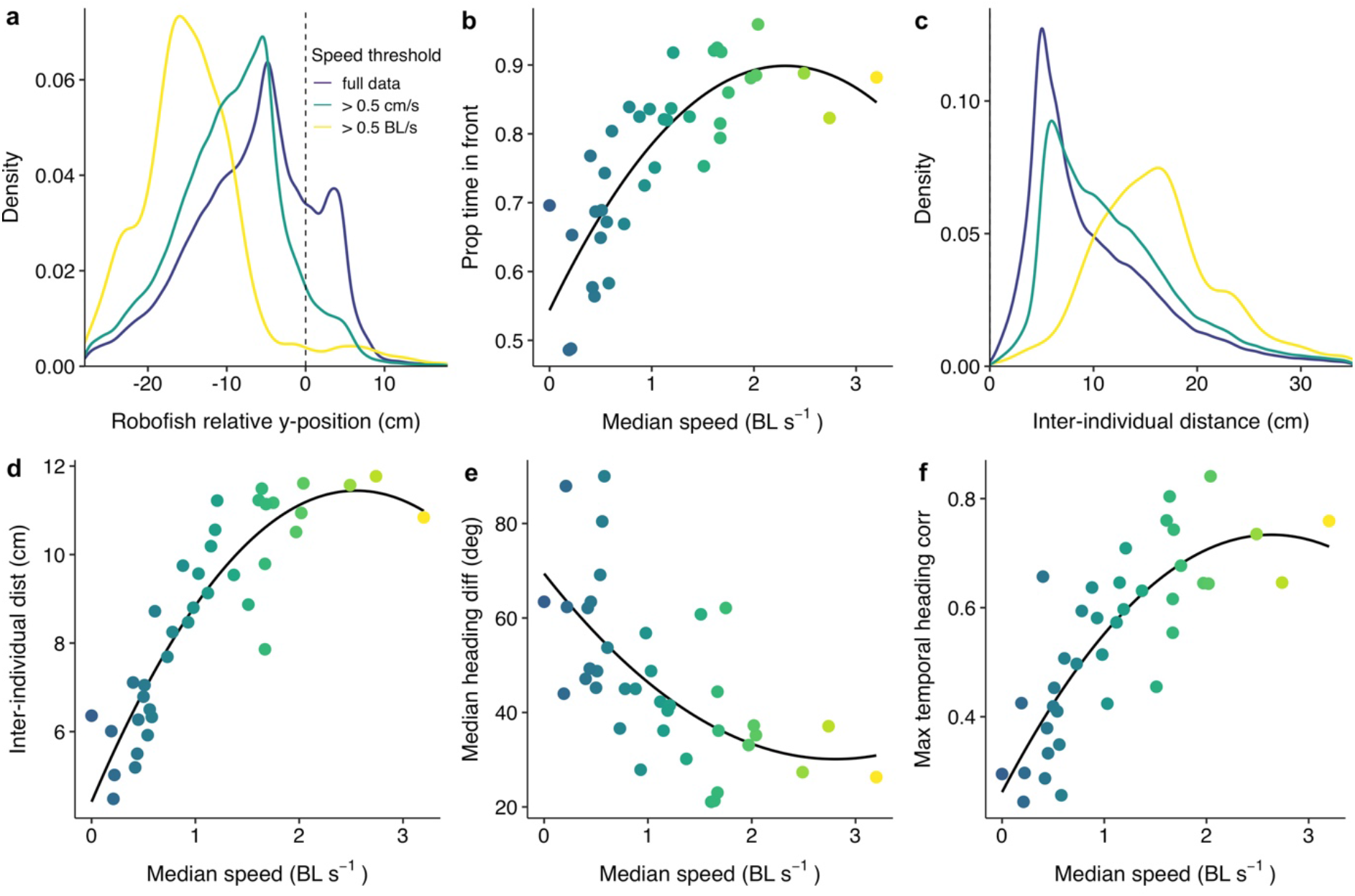
(a,c) Density plot of robofish’s relative y-position and distance to their live partner, and (b,d,e,f) scatterplots of fish’s median speed in relation to the leadership, cohesion, alignment, and temporal coordination of the pair. Colour scale indicates speed (blue=low; yellow=high) and solid lines show the polynomial functions fitted in our models.

Robofish was able to maintain good cohesion and alignment with its live partner, and naturally copied its changes in speed and heading (median max correlation coefficients: 0.51 and 0.58, Figure S7). As for leadership we found that these group-level outcomes were very well explained by the individual speed of the guppy. Pairs in which the guppy had a high median speed were considerably less cohesive (χ^2^ = 60.04, *p* < 0.001, R^2^_mar_ = 0.81), more aligned (χ^2^ = 13.66, *p* < 0.001, R^2^_mar_ = 0.25), and more coordinated (χ^2^ = 41.33, *p* < 0.001, R^2^_mar_ = 0.65) than pairs in which the guppy was much slower (Figure 2c-f).

## Discussion

Live guppies paired with an extremely social robotic fish showed large and repeatable individual differences in movement speeds that in turn strongly explained leadership, group cohesion, alignment, and movement coordination. By testing all fish with a robot that used identical interaction rules and lacked any preferred movement speed and directionality, these results provide novel experimental evidence that suggests individual speed is a fundamental factor in the emergence of collective behavioural patterns, in line with existing theoretical and empirical work [4,8,12,13,23]. As individual differences in speed are associated with a broad range of phenotypic traits observed among grouping animals, this may also help provide a mechanistic explanation for the effect of phenotypic heterogeneity for group-level patterns [3], such as has been shown for size, hunger, and parasitism [24,25].

We observed a very strong positive link between speed and leadership, both in terms of clearer front-back positioning the faster fish were moving, as well as fish being overall more in front the higher their median movement speed. This result is in line with predictions from model simulations [4,12] and previous observational studies [11,12]. By testing fish with an extremely social partner that always tried to follow, we found that at higher speeds fish led almost all the time. This shows how leaders depend on the responsiveness of their followers in order to express their own preference (see also [26]) and more generally highlights how both individual speeds and high levels of social responsiveness are important for the collective performance of groups [27]. At lower speeds leadership differences were not as apparent. This could potentially be explained by individuals’ increased freedom for turning at lower speeds [8] and indicates that besides relative differences in speed between group members also their absolute speed is important in shaping group structure.

Pairs consisting of a guppy with a high median speed were considerably less cohesive than those with low median speeds, in line with previous work on schooling sticklebacks [12]. To avoid collisions, grouping animals may actively increase their distance when moving at higher speeds. However, robofish was not programmed with such a rule, suggesting that the observed positive link between cohesion and speed is due to speed mediating the use of social interaction rules: faster individuals moving farther before they can change their position based on their group mates. This suggests that a shift in interaction rules, such as by changes in the environment, may alter the relationship between group speed and cohesion (see e.g. [28,29]). Individual speed also strongly drove the alignment and temporal coordination of the pairs, in line with previous empirical studies that found fast moving groups tend to be polarised and slow moving groups to be disordered [12,13,30]. As in our study the robotic partner completely lacked any alignment rules, our findings provide novel empirical evidence that individual speed is a key factor facilitating group alignment and coordination. Although speed-mediated changes in local interaction rules may help to explain these effects [6,10], groups may be more likely to become disordered at lower speeds because of larger potential angular fluctuations at lower speeds, as is predicted from theoretical analysis [8]. Our finding that faster groups showed better coordination of movement changes and information flow being higher in faster groups, as shown by previous work [13], can thereby be directly explained by the higher (local) order that arises with higher individual speeds.

The large individual differences in movement speed during the robofish trials were highly repeatable (R = 0.70), as compared to the average repeatability of 0.37 reported by a large meta-analysis [31]. As robofish always copied its live partner’s speed and movements and always used the same interaction rules, the large variability in movement speed among the pairs must be attributable to the speed of the live fish, which was well explained by fish’s solo speed. Interestingly however, considerable speed variation among the fish remained. The reduction in movement speed between the solo and robofish trials therefore likely reflects socially-mediated changes, with guppies that slowed down more being more socially responsive and/or less inclined to lead. This corroborates previous observational work that found live fish pairs moved faster when led by fish that were less socially plastic in their speeding changes [7]. Future work is needed to properly determine to what extent this behavioural variation in ‘social speed’ indicates true individual differences in social responsiveness.

In summary, by closed-loop experiments of live guppies swimming with a biomimetic robot that always followed and naturally copied its partners’ movements, we provide novel evidence that individual speed is a fundamental factor for the emergence of collective behaviour. By programming the robotic fish without any of its own movement preferences, we had the unique opportunity to investigate how individual behaviour leads to group-level patterns without the potential influence of individual heterogeneity in group mates. Exciting interdisciplinary work lies ahead to further investigate the role that individuals play in animal groups and how that depends on the social feedback among heterogeneous group members.

## Supporting information

Summary data

## Acknowledgements

We acknowledge financial support from the Alexander von Humboldt-Stiftung (postdoctoral fellowship to JWJ), the Zukunftskolleg, University of Konstanz (postdoctoral fellowship to JWJ), the German Research Foundation (BI 1828/2-1, LA 3534/1-1, RO 4766/2-1) and Germany’s Excellence Strategy (EXC 2002/1 “Science of Intelligence”, project number 228 390523135).

## Authors’ contributions

J.W.J, N.W. and D.B. designed the study, T.L., D.B, P.R. and J.K. developed robofish, N.W. and D.B conducted the experiment, J.W.J. performed the analysis, J.W.J and D.B. wrote the manuscript with feedback from all other authors.

