## Supplementary methods and figures for "Group-level patterns emerge from individual speed as revealed by an extremely social robotic fish"

For:

### Appendix A: Supplementary Methods

#### Experiments

For our experiments we used only females to avoid sex differences in attraction toward Robofish, which resembles a female guppy. Fish were tested in 7.5cm of water to allow the replica to move close to the surface while simultaneously avoiding surface disturbances that might interfere with tracking. Robofish trials started with the acclimatisation cylinder being raised and the robofish performing a circular movement with a diameter of 20 cm at a speed of 8 cm/s in front of the holding cylinder to attract the focal fish. As soon as the fish moved, the milling behaviour was terminated and robofish was initiated to move towards the live guppy. The 3D-printed replica we used for our robotic experiments was designed to resemble a female guppy and is readily followed by live fish because it has key features such as glass eyes and natural motion patterns [1]. At its base it has a magnet that aligns with a second magnet below the tank that sits on top of a two-wheeled robot unit that can move freely on a second level below the tank, thus coupling the robot's motions directly with the replica (see Figure S1). By using a biomimetic robotic fish, we were able to create an extremely social partner without any personal movement preferences that showed natural social interactions. This is clear from the extremely high correlation in movement speeds between the robotic fish and its live partner and the observation of overall highly cohesive, well-coordinated movements. As in previous studies [2], robofish seem to be accepted as a conspecific by our live test fish, which is underpinned by the fact that a majority of fish reduced their movement speed when interacting with robofish and did not show a flight reaction towards the constantly approaching robot (see Figure S6b).

#### Data processing

Standard lengths of the fish were determined by taking a picture of each fish in a transparent water container positioned on graph paper, and the length from the tip of fishes' snout to the caudal base determined using ImageJ 1.52. We used BioTracker [3] to automatically acquire detailed coordinate data (centre of mass) of the fish. Two robotic trials had to be excluded due to issues with recording. Trajectories were carefully checked for any errors and the first 175 frames of each trial removed to account for the closed-loop system requiring time to establish itself. Coordinates were subsequently converted from pixels to mm. From the tracking data, we computed each individual's velocity and heading, and their speed based on a moving window of 5 frames to account for short, spurious changes in speed. Speed was subsequently computed in terms of standard body lengths to account for any potential body size effects. For the robofish trials, we additionally computed each pairs' cohesion, in terms of the distance between the live guppy and the robofish, and alignment, in terms of the absolute difference in heading. We also computed fishes' front-back positioning to determine which individual in the mixed pairs was more likely to lead. We shifted the coordinates of the focal fish such that its origin was at the origin pointing north, and counted the proportion of frames that the robofish had a negative relative y-position (i.e. behind the live fish). Finally, to determine the propagation of movement changes in the pairs, we ran temporal correlations on their speed and heading, with a maximum delay of 5 seconds, and determined the average directional correlation between the two fish as a function of the delay in time. For each trial we computed fishes' median speed, median heading difference, median inter-individual distance, proportion of time the focal fish was in front, and the max temporal correlation coefficients across all frames.

**Data analysis**

We used a generalised linear mixed modelling (LMM) approach to investigate the repeatability in individual speed as well as the link between individual speed to our measures of interest. Specifically, to compute behavioural repeatability, we ran mixed models with fishes' median speed as response variable and individual ID as a random factor, and ran 10,000 permutations to acquire 95% confidence intervals, which indicate a significant effect when there is no overlap with 0 [4]. To investigate how fishes' speed during the robofish trials was related to their solo speed, we ran a model with fish' median speed as response variable, fish' solo speed as a fixed factor, and their ID as a random factor. To investigate the correlation in speed between the robofish and their live partner, we ran a model with the median speed of the robofish as response, their live partner's median speed as a fixed factor, and fish ID as a random factor.

To investigate how guppies' median speed was linked to leadership, cohesion, alignment, and temporal coordination (e.g., how well turning changes propagated within the pairs), we ran separate models with respectively proportion of time the guppy was in front, fishes' median inter-individual distance, fishes median heading difference, and median temporal heading coordination as response variables, fishes' median speed during the robo trials as a fixed effect, and individual ID as a random factor. We included orthogonal polynomials, as a quadratic term significantly improved movel fit compared to the linear term (all final models $\Delta AIC < 2$), for which we used the poly() function in R to reduce multicollinearity. Body size was also included as a fixed factor in all models but never found to be significant and not described further. For our LMM approach, models were fitted with a Gaussian error distribution. Minimal adequate models were obtained using backward stepwise elimination. Residuals were visually inspected to ensure homogeneity of variance, normality of error, and linearity. As heading changes are increasingly spurious at lower speeds, we also ran a model with group alignment computed for the data subsetted to cases when the live guppy moved at a speed of at least 0.5 cm/sec, which did change the model fit. Means are quoted ± SE unless stated otherwise. All data were analysed in R 3.5.0.

1. Landgraf T, Bierbach D, Nguyen H, Muggelberg N, Romanczuk P, Krause J. 2016 RoboFish: increased acceptance of interactive robotic fish with realistic eyes and natural motion patterns by live Trinidadian guppies. *Bioinspir. Biomim.* **11**, 015001. (doi:10.1088/1748-3190/11/1/015001)
2. Bierbach D, Landgraf T, Romanczuk P, Lukas J, Nguyen H, Wolf M, Krause J. 2018

Using a robotic fish to investigate individual differences in social responsiveness in the guppy. *R. Soc. Open Sci.* **5**, 181026. (doi:10.1098/rsos.181026)
3. Mönck HJ *et al.* 2018 BioTracker: An Open-Source Computer Vision Framework for Visual Animal Tracking. *arxiv*: 1803.07985.
4. Dingemanse NJ, Dochtermann NA. 2013 Quantifying individual variation in behaviour: mixed-effect modelling approaches. *J. Anim. Ecol.* **82**, 39–54. (doi:10.1111/1365-2656.12013)

### Appendix B: Supplementary Figures

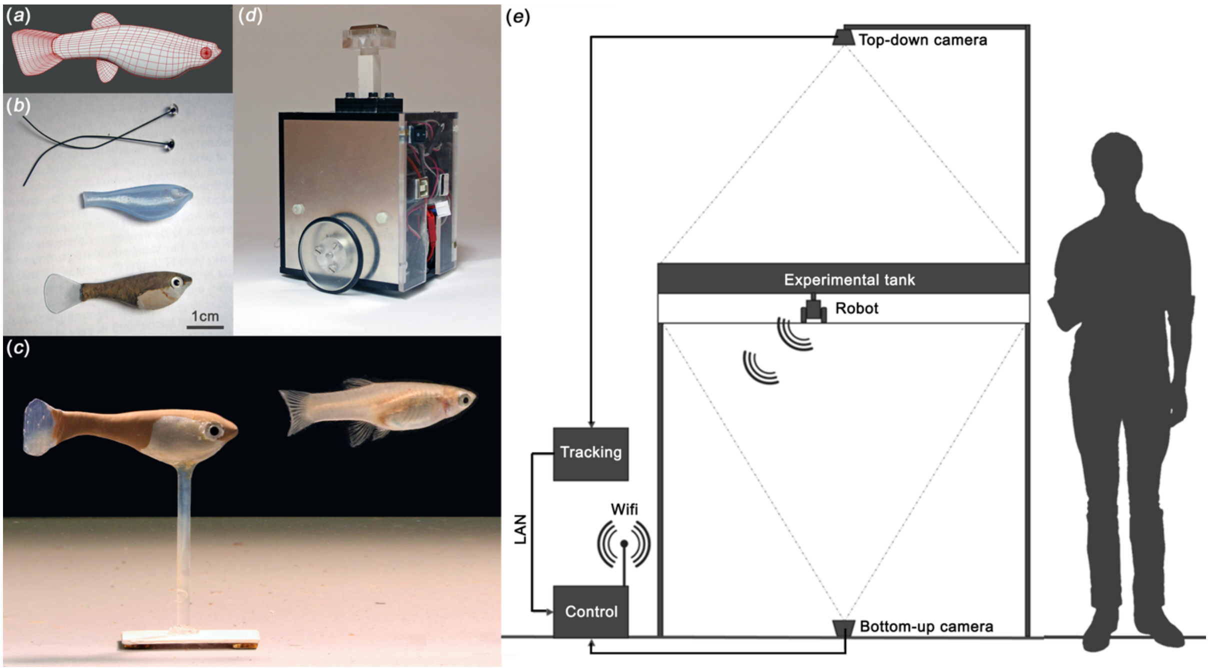


**Figure S1.** (a) The 3D-mesh that we used to print our replicas, which was based on photographs of a live adult female; (b) the guppy-like replica equipped with glass eyes and transparent caudal fin to resemble a live guppy female; (c) the replica fixed on its transparent plastic stick, glued to a plastic plate containing the magnet; (d) close-up picture of the two-wheeled robot unit that was used to move the replica in the tank; (e) experimental setup showing the test tank and cameras above and below the tank as well as their connectivity.

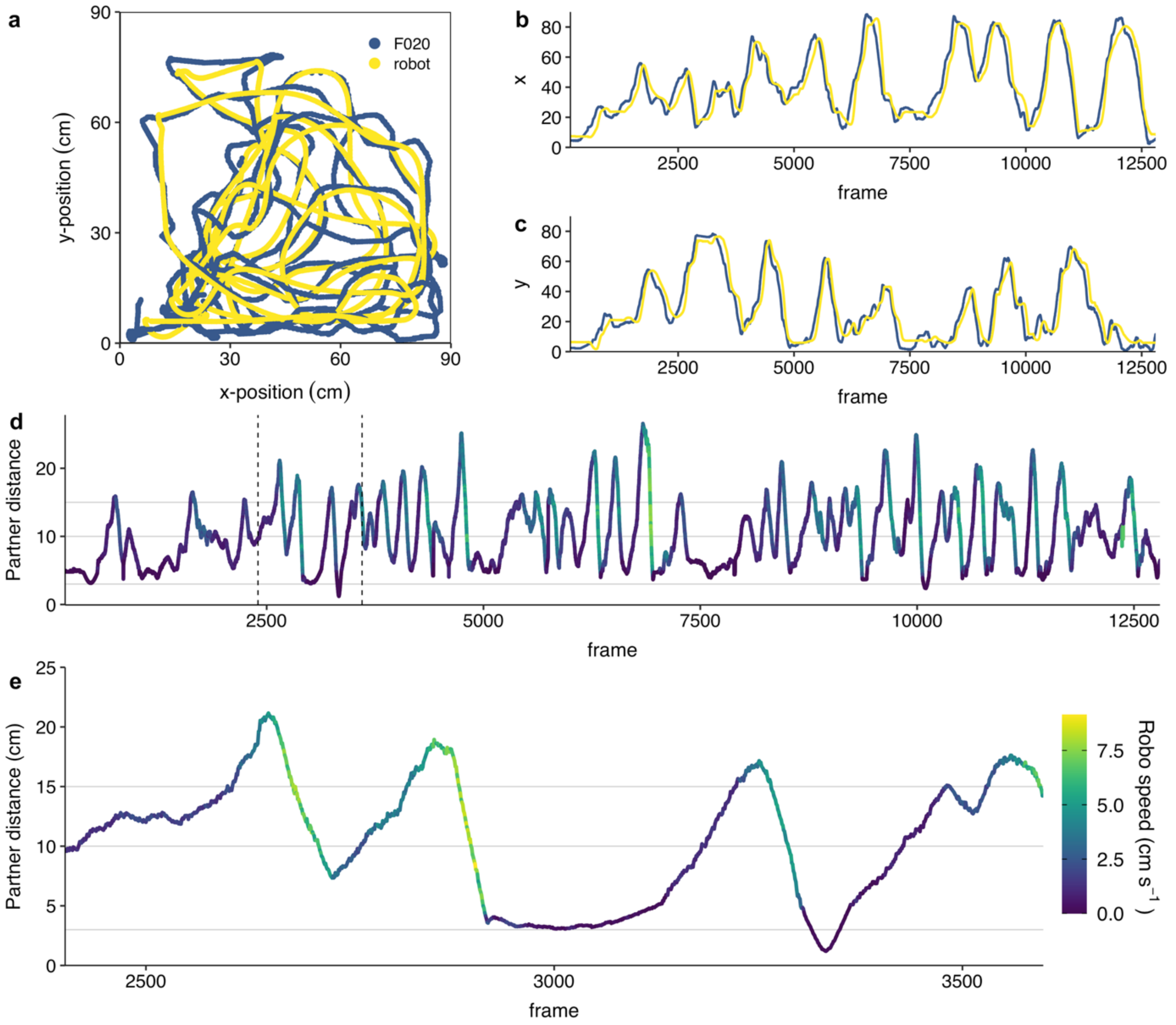


**Figure S2.** Detailed spatial and temporal plots of the tracking data, cohesion, and speed of a randomly selected robofish trial. (a-c) Trajectories of the live guppy (depicted in blue) and the robofish (in yellow) in two-dimensions (a), and one-dimensional for the x- and y-coordinate (b,c). (d,e) Detailed temporal plots of the link between the inter-individual distance of robofish relative to its partner and its speed (blue=low; yellow=high). Plots (a-d) show the full-frame-by-frame data of one randomly selected trial, while plot (e) shows a subset of the same trial (~ 1min), depicted by the dashed lines in plot (c), that contains some key changes in speed. These plots show the robofish was able to stay very cohesive with its partner and carefully followed and copied its movements, as indicated by the trajectories being close together and with a slight temporal delay for the robot. Furthermore, the robofish was able to stay close to its partner when it swam further away by significant increases in speed, and avoided getting too near by slowing down completely. Figure 1 in the main text is based on the same datafile and subsetted trajectory data as that depicted here.

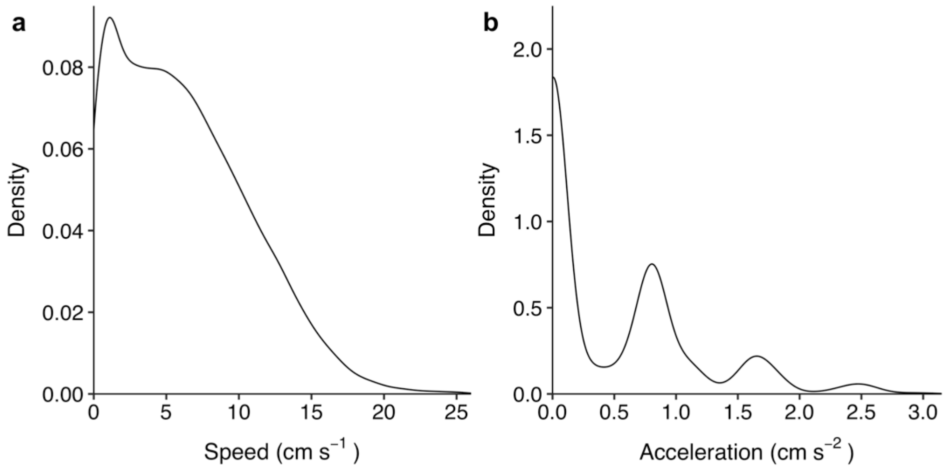


**Figure S3.** Density plots of guppies' speed (a) and acceleration (b) during the solo trials ($n$ = 20 fish), based on the frame-by-frame data, with both measures being computed with a rolling window of 3fps to control for very short spurious changes in speed.

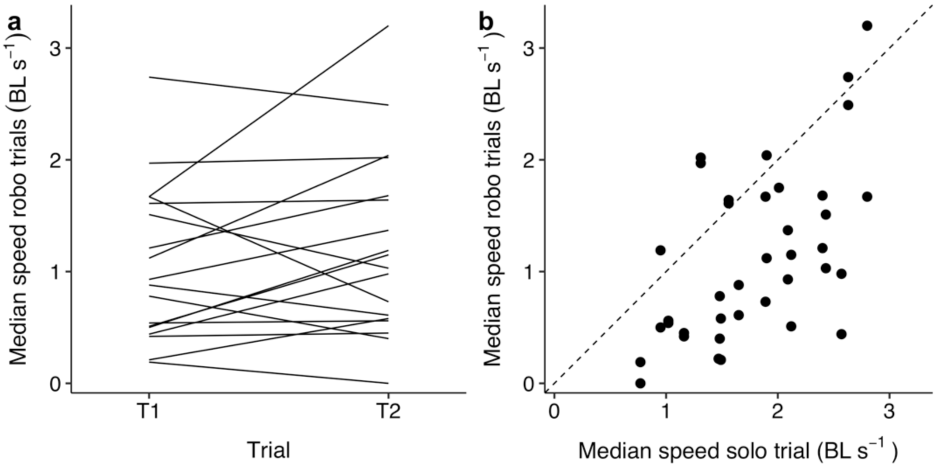


**Figure S4.** (a) Line plot of guppies' median speed during the robofish trials ($n$ = 38 trials); (b) scatter plot of guppies' median speed during the robofish trials as a function of their speed during the solo trial ($n$ = 20 fish). These plots show there was large among-individual variation in movement speeds during the robofish trials, with fish being highly repeatable across the trials and speed being strongly linked to their solo speed.

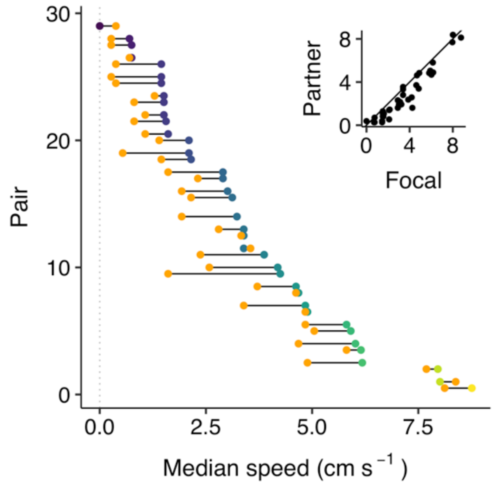


**Figure S5.** Plot showing the median speed of each fish (dots coloured blue to yellow for low to high speeds) and its robotic partner (orange dots) for the 38 robofish trials. Inset shows the same data but with the speed of the live guppy and the robofish projected on the x and y axes respectively. This plot shows there was large variation among the fish but high conformity in speed within the guppy-robofish pairs.

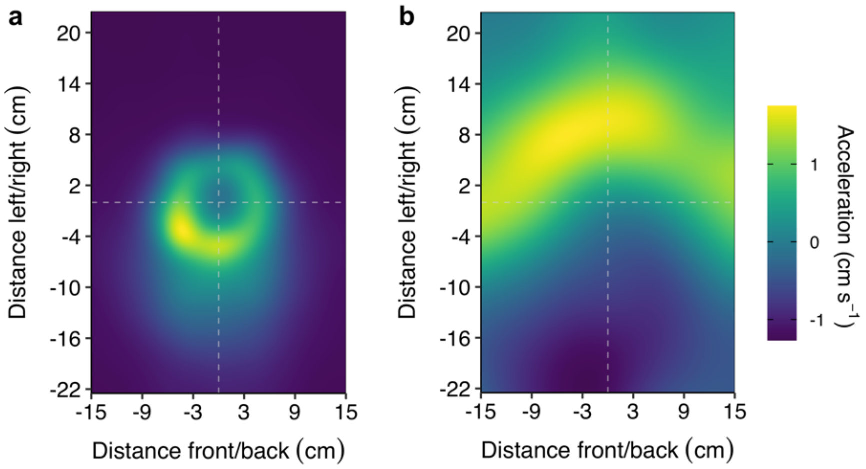


**Figure S6.** (a) Heatmap of the spatial positioning of the robofish relative to the position of the focal fish at the origin pointing north, based on the frame-by-frame data of all robofish trials (*n* = 38). (b) Heatmap of guppies' acceleration as a function of the relative position of the robofish. For both plots data was cropped to 93% of the full parameter space to show the most relevant area only. The colour scale is proportional to the densest bin of each plot (blue = low; yellow = high). Plot (a) shows that the robofish was primarily behind the focal fish and rarely in a zone roughly equating the repulsion zone of 3cm it was programmed to avoid. Plot (b) shows fish sped up in the rare cases when the robofish was in front, and slowed down when it was behind, but at a relatively much further distance. This indicates the focal fish also responds to the position of the robofish, but is not that socially responsive to it to maintain cohesion when it is far behind.

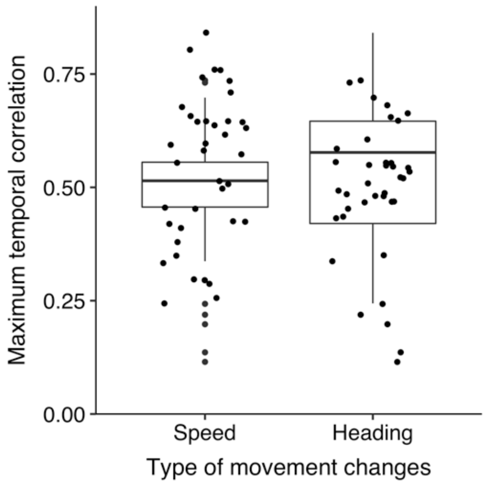


**Figure S7.** Boxplots of the maximum temporal correlations of speed and heading changes between the robofish and its live partner for the robofish trials ($n$ = 38). For all trials, the maximum correlation occurred with the data of the robofish shifted with a delay relative to the focal fish, rather than the other way around, indicating the robofish primarily copied the movement changes of the live fish.
